# A novel systems biology approach to evaluate mouse models of late-onset Alzheimer’s disease

**DOI:** 10.1101/682856

**Authors:** Christoph Preuss, Ravi Pandey, Erin Piazza, Alexander Fine, Asli Uyar, Thanneer Perumal, Dylan Garceau, Kevin P Kotredes, Harriet Williams, Lara M Mangravite, Bruce T. Lamb, Adrian L. Oblak, Gareth R. Howell, Michael Sasner, Benjamin A Logsdon, Gregory W. Carter

## Abstract

**Background:** Late-onset Alzheimer’s disease (LOAD) is the most common form of dementia worldwide. To date, animal models of Alzheimer’s have focused on rare familial mutations, due to a lack of frank neuropathology from models based on common disease genes. Recent multi-cohort studies of postmortem human brain transcriptomes have identified a set of 30 gene co-expression modules associated with LOAD, providing a molecular catalog of relevant endophenotypes.

**Results:** This resource enables precise gene-based alignment between new animal models and human molecular signatures of disease. Here, we describe a new resource to efficiently screen mouse models for LOAD relevance. A new NanoString nCounter® Mouse AD panel was designed to correlate key human disease processes and pathways with mRNA from mouse brains. Analysis of three mouse models based on LOAD genetics, carrying APOE4 and TREM2*R47H alleles, demonstrated overlaps with distinct human AD modules that, in turn, are functionally enriched in key disease-associated pathways. Comprehensive comparison with full transcriptome data from same-sample RNA-Seq shows strong correlation between gene expression changes independent of experimental platform.

**Conclusions:** Taken together, we show that the nCounter Mouse AD panel offers a rapid, cost-effective and highly reproducible approach to assess disease relevance of potential LOAD mouse models.

## BACKGROUND

Late-onset Alzheimer’s disease (LOAD) is the most common cause of dementia worldwide (1). LOAD presents as a heterogenous disease with highly variable outcomes. Recent efforts have been made to molecularly characterize LOAD using large cohorts of post-mortem human brain transcriptomic data (2). Systems-level analysis of these large human data sets has revealed key drivers and molecular pathways that reflect specific changes resulting from disease (2,3). These studies have been primarily driven by gene co-expression analyses that reduce transcriptomes to modules representing specific disease processes or cell types across heterogenous tissue samples (2,4,5). Similar approaches have been used to characterize mouse models of neurodegenerative disease (6). Detailed cross-species analysis reveals a translational gap between animal models and human disease, as no existing models fully recapitulate pathologies associated with LOAD (7,8). New platforms to rapidly assess the translational relevance of new animal models of LOAD will allow efficient identification of the most promising preclinical models.

In this study, we describe a novel gene expression panel to assess LOAD-relevance of mouse models based on expression of key genes in the brain. We used a recent human molecular disease catalog based on harmonized co-expression data from three independent post mortem brain cohorts (ROSMAP, Mayo, Mount Sinai Brain bank) (9–11) and seven brain regions that define 30 human co-expression modules and five consensus clusters derived from the overlap of those modules (12). These modules were used to design a mouse gene expression panel to assess the molecular overlap between human disease states and mouse models. This nCounter Mouse AD panel was piloted with samples from three novel mouse models of LOAD. Same-sample comparison between NanoString and RNA-Seq data demonstrated high per-gene correlation and overall concordance when separately compared to human disease co-expression modules. Taken together, the rapid screening of mouse models in the course of different life stages will allow better characterization of models based on alignment with specific human molecular pathologies.

## RESULTS

### Human-mouse co-expression module conservation and probe coverage across 30 LOAD associated modules

An overview of the Mouse AD panel design for translating the 30 human AMP-AD co-expression modules from three cohorts and seven brain regions is depicted in Figure 1. Mouse to human gene prioritization resulted in the selection of 760 key mouse genes targeting a subset of highly co-expressed human genes plus 10 housekeeping genes, which explained a significant proportion of the observed variance across the 30 human AMP-AD modules (Methods). Co-expression modules were grouped into functionally distinct consensus clusters as previously described by Logsdon, et al (see also Supplemental Table 1) (12). These consensus clusters contain expression modules from different brain regions and independent studies that share a high overlap in gene content and similar expression characteristics. Consensus clusters were annotated based on Reactome pathway enrichment analysis for the corresponding genes within each functionally distinct cluster (Methods, Supplemental Table 1). Since consensus clusters showed an enrichment of multiple biological pathways, the highest rank and non-overlapping Reactome pathway was used to refer to each cluster (Supplemental Table 2). In order to assess the conservation of sequence and gene expression levels between human and mouse genes for each of the 30 human co-expression modules, dN/dS values were correlated with the overall overlap in expression in brains from six-month-old C57BL/6J (B6) mice (Figure 2A). The fraction of orthologous genes expressed in the mouse brain, based on the presence or absence of transcripts at detectable levels, was very highly correlated with the overall module conservation (p-value < 2.2e-16, Pearson’s correlation coefficient: −0.96). Module conservation was based on the median dN/dS statistics measuring the rate of divergence in the coding sequence for all genes within a given module between both species (Figure S1). Notably, human co-expression modules of Consensus Cluster C, associated with the neuronal system and neurotransmission, showed the lowest degree of sequence divergence with a high proportion of human genes (64-72%) expressed in six-month-old B6 mice. In contrast to the highly conserved neuronal modules, immune modules of Consensus Cluster B contained genes that recently diverged on the sequence level and acquired a higher number of destabilizing missense variants. These modules showed the highest median dN/dS values and the lowest fraction of genes (27-46%) expressed in the mouse brain across all tested modules. The remaining human co-expression modules, associated with different functional categories (Figure 2A, Supplemental Table 1), had intermediate overlap in expression levels between human and mice. Each of the 30 human co-expression modules was covered with an average of 148 NanoString mouse probes (SD = 50 probes), where a single mouse probe can map to multiple human modules from different study cohorts and across several brain regions. Overall, mouse probe coverage for human co-expression modules ranged between 4% and 19%, depending on the size and level of conservation of the targeted human module (Figures 2B and 2C, Supplemental Tables S2 and S3). For three of the largest human co-expression modules harboring over 4,000 transcripts, the probe coverage was slightly below the targeted 5% coverage threshold. However, these large modules are predominantly associated with neuronal function and show a high degree of expression and sequence conservation between human and mouse (Figures 2A). Immune modules, containing genes that recently diverged on the coding sequence level, are well covered with a median coverage of 10% (Figure 2C). A complete annotation of mouse probes to human transcripts for each human co-expression module is provided in Supplemental Table S3. In addition, we compared our novel panel to the existing nCounter Mouse Neuropathology panel designed to assess expression changes in multiple neurodegenerative diseases. We observed an overlap of 105 probes (7%) between both panels, highlighting that most of our selected probe content is novel and specific to LOAD associated disease processes and pathways.

**Figure 1:**
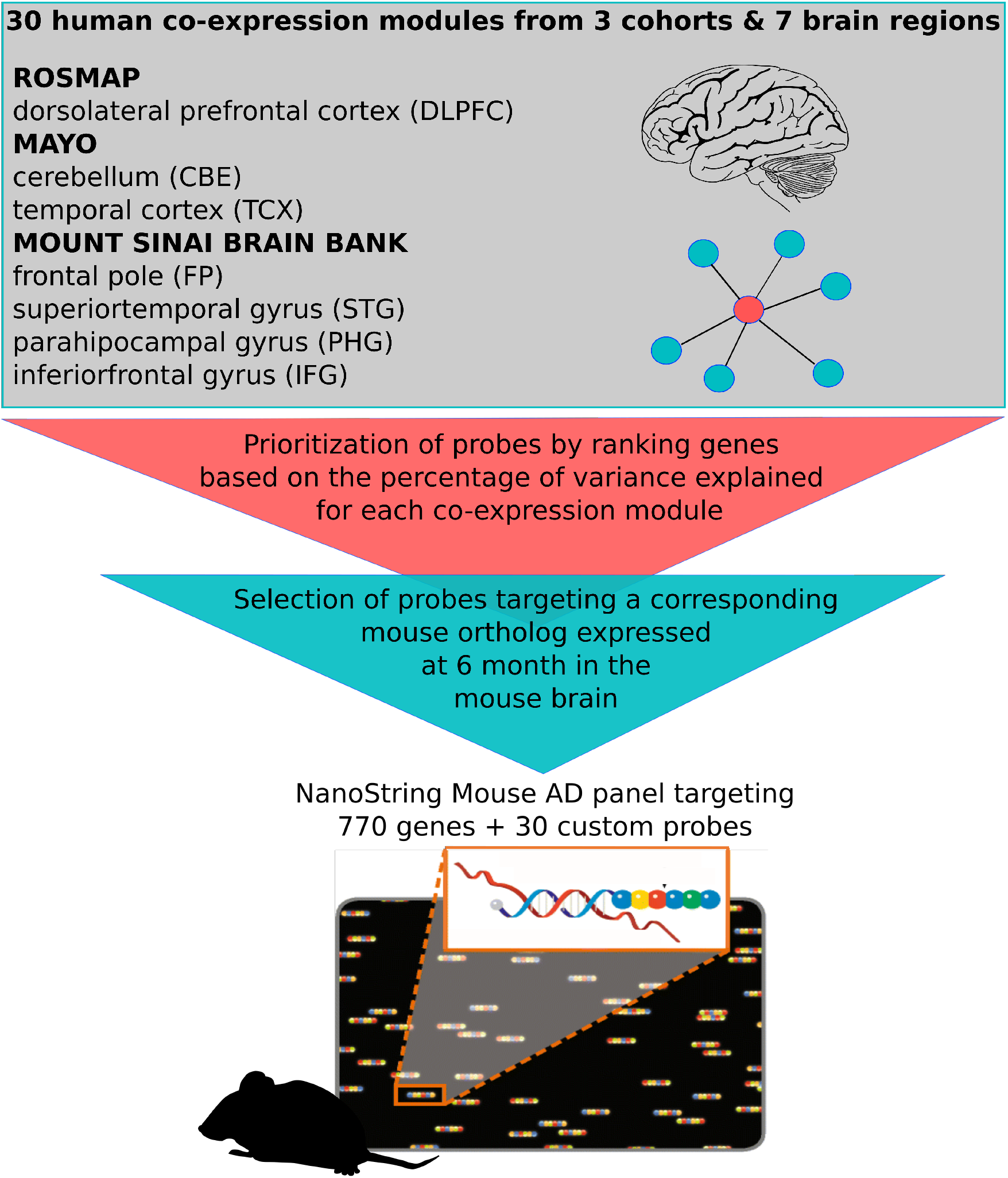
Overview of the nCounter Mouse AD panel design. The novel Mouse AD panel measures expression of genes from a set of 30 human co-expression modules from three human LOAD cohorts, including seven distinct brain regions. Human genes central to each of the human expression modules were prioritized for the Mouse AD panel to select conserved signatures of LOAD associated pathways.

**Figure 2:**
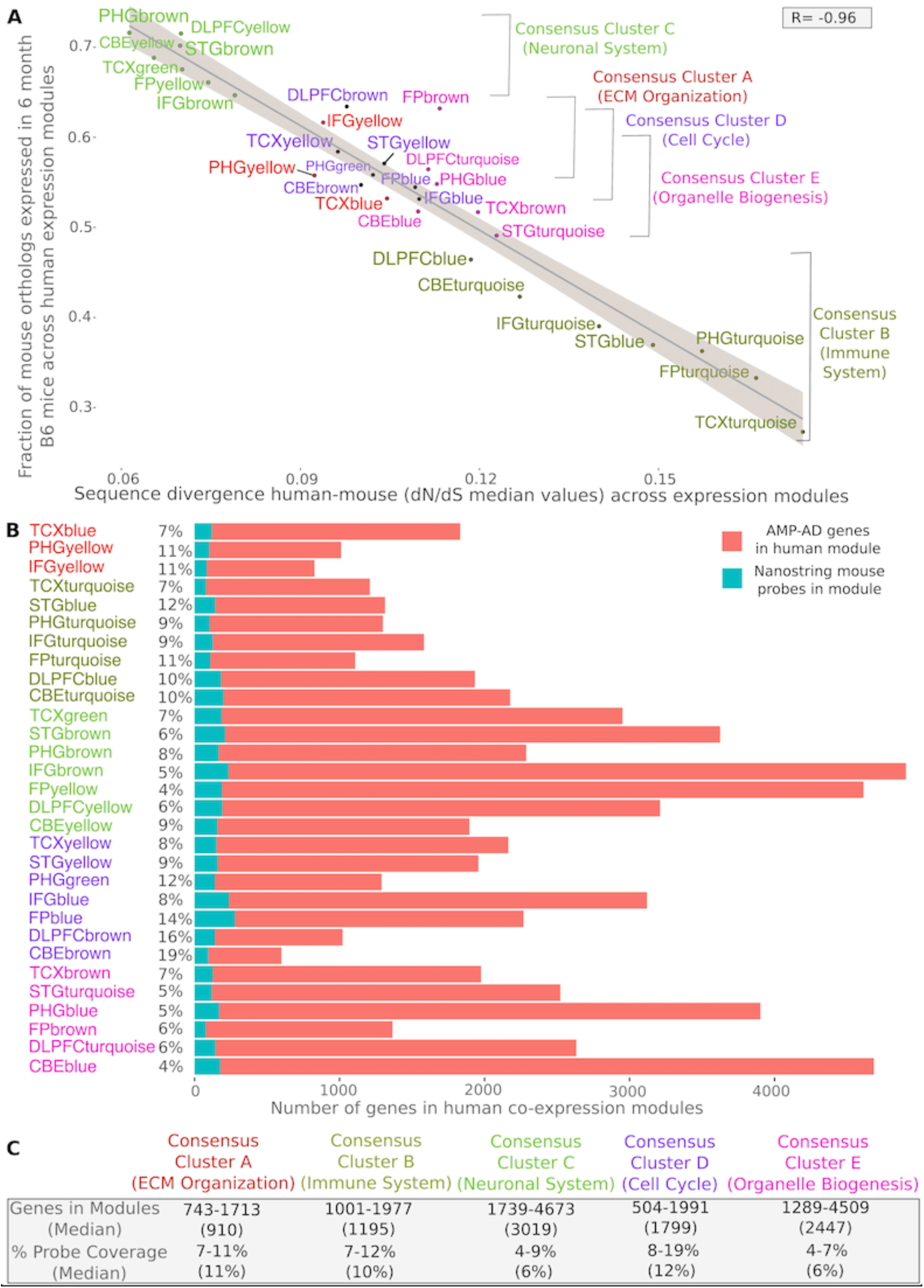
Human to mouse comparison and probe coverage summary statistics. **(A)** Human-mouse sequence divergence (median dN/dS values) is inversely correlated (Pearson’s correlation coefficient: −0.96) with the fraction of genes being expressed in B6 mouse brain for each of the human co-expression modules. **(B)** Coverage of the 770 selected mouse NanoString probes for the 30 human co-expression modules associated with five functional consensus clusters. The size and number of human co-expression modules differs for the three post-mortem brain cohorts (ROSMAP, Mayo, Mount Sinai Brain Bank) and across the seven included brain regions. (**C**) This results in a varying degree of probe coverage for each module with a number of disease associated consensus clusters (A-E), reflecting disease related pathways and processes.

### Prioritized subset of key genes show a higher degree of sequence conservation and expression level across modules

In order to assess the level of sequence divergence and expression for the prioritized subset of genes on the novel panel, the selected subset of genes were compared to all genes across the 30 human co-expression modules. The 760 key genes, explaining a significant proportion of the observed variance in each human module, showed an overall lower level of sequence divergence (median dN/dS values) when compared to all other genes in the modules (Figure 3, Figure S1). Furthermore, the selected key genes on the Mouse AD panel also displayed a higher average level of gene expression in brains of six-month-old B6 mice compared to the remaining genes for each of the 30 modules (Figure 3). This highlights that our formal prioritization procedure resulted in the selection of a subset of highly expressed key genes, which are also more conserved between human and mouse facilitating the translation of co-expression profiles across species.

**Figure 3:**
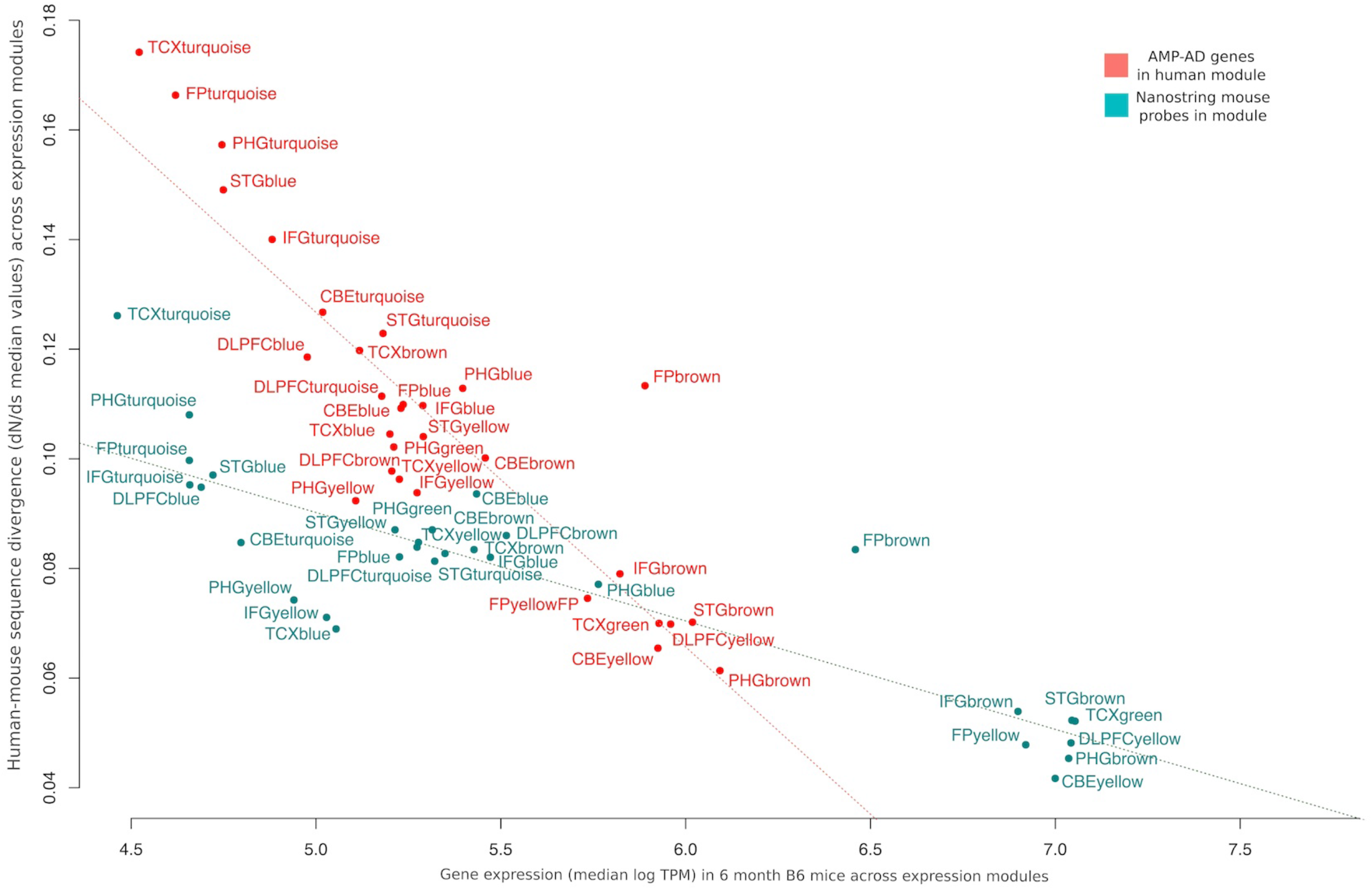
NanoString Mouse AD probe genes are more conserved and have greater expression in the mouse brain. Comparison between gene-level sequence divergence and transcript abundances in six month old B6 mouse brains for all genes (red) and the subset of 770 genes covered by NanoString probes (green).

### Novel mouse models harboring LOAD associated risk variants correlate with distinct AMP-AD modules in a brain region- and pathway-specific manner

Three novel mouse models, harboring two LOAD risk alleles, (Supplemental Table S4) were used to translate co-expression profiles between human and mouse brain transcriptome data using our novel Mouse AD panel. Transcriptome analysis was performed for the APOE4 KI mouse, carrying a humanized version of the strongest LOAD associated risk allele (*APOE-ε4*) and the Trem2*R47H mouse, which harbors a rare deleterious variant in *TREM2*. The rare *TREM2* R47H missense variant (rs75932628) has been previously associated with LOAD in multiple independent studies [16,17]. In addition, a mouse model harboring both, the common and rare AD risk variants (APOE4 KI/Trem2*R47H) was used to compare the transcriptional effects in mice carrying both variants to mice carrying only a single risk allele and B6 controls. Mouse transcriptome data for half brains was analyzed at different ages (4-14 months) to estimate the overlap with human post-mortem co-expression modules during aging. We observed specific overlaps with distinct disease processes and molecular pathways at different ages for the APOE4 KI and Trem2*R47H mouse models. At an early age (2-5 months), male APOE4 KI and Trem2*R47H mice showed strong positive correlations (p-value < 0.05, Pearson’s correlation coefficient < −0.3) with human co-expression modules in Consensus Cluster E that are enriched for transcripts associated with cell cycle and RNA non-mediated decay pathways in multiple brain regions (Figure 4). Furthermore, Trem2*R47H male mice showed a significantly negative association (p-value < 0.05, Pearson’s correlation coefficient < −0.2) with immune related human modules in the superiortemporal gyrus, the inferiorfrontal gyrus, cerebellum and prefrontal cortex (Figure 4). This effect becomes more pronounced later in development, between six and 14 months, when the correlation with human immune modules is also observed in Trem2*R47H female mice. During mid-life, (6-9 month-old age group), we observed an age-dependent effect for the APOE4 KI mouse in which human neuronal modules in Consensus Cluster C start to become positively correlated with the corresponding human expression modules (Figure 4). Interestingly, neuronal co-expression modules which are associated with synaptic signaling appear to be positively correlated with APOE4 KI, but not Trem2*R47H mice in an age dependent manner. This up-regulation of genes associated with synaptic signaling and a decrease of transcripts enriched for cell cycle, RNA non-mediated decay, myelination and glial development in aged mice was consistent for multiple brain regions and across three independent human AD cohorts. When compared to APOE4 KI mice, Trem2*R47H mice showed an age dependent decrease in genes associated with the immune response in several AMP-AD modules which is not observed for APOE4 KI mice (Figure 4). Notably, the APOE4 KI/Trem2*R47H mice showed characteristics of both single variant mouse models. At an early age, an overlap with both neuronal and immune associated human modules is observed and becomes more pronounced during aging.

**Figure 4:**
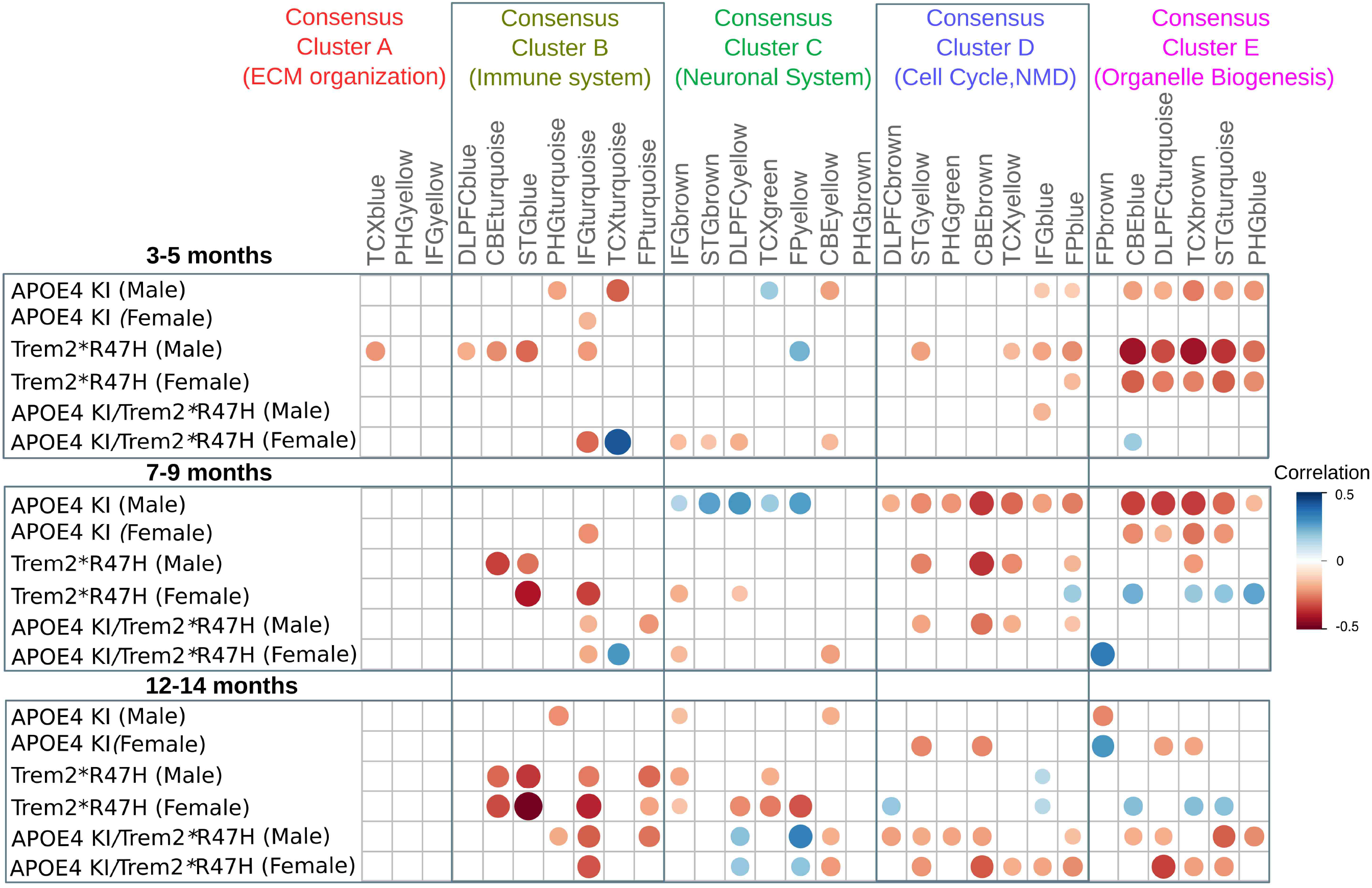
Correlations between mouse model effects and LOAD effects for the NanoString Mouse AD panel genes across 30 human co-expression modules. Circles correspond to significant (p-value < 0.05) positive (blue) and negative (red) Pearson’s correlation coefficients for gene expression changes in mice (log fold change of strain minus age-matched B6 mice) and human disease (log fold change for cases minus controls). Human co-expression modules are ordered into Consensus Clusters A-E (12) describing major sources of AD-related alterations in transcriptional states across independent studies and brain regions. Consensus clusters are annotated based on the most significantly enriched and non-redundant Reactome pathway terms (Supplemental Tables S1, S2).

### Comparison between nCounter Mouse AD panel and RNA-Seq data

To assess the validity of the novel Mouse AD panel across transcriptomic platforms, we compared the results from the nCounter analysis to results from RNA-Seq data for the same 137 mouse brain samples. A correlation analysis was performed to compare the expression of the 770 NanoString probes across co-expression modules with RNA-Seq transcript expression for all ages (3-5, 7-9, 12-14 months), highlighting the different LOAD mouse models as independent variables (Figure 5). For the direct comparison, between the 770 NanoString probes with corresponding RNA-Seq transcripts, a similar range of correlation coefficients between human data and the three mouse models was observed (Figure 5A). Overall, the correlation between the RNA-Seq and NanoString platforms were high across all age groups (Pearson’s correlation coefficients: 0.65-0.69) when comparing the subset of 760 key transcripts and 10 housekeeping transcripts across platforms. This demonstrates that the novel NanoString panel, despite the limited number of key custom probes, can achieve similar results when compared to high-throughput RNA-Seq data. Furthermore, the alignment of human and mouse modules based on the expression of all genes within each modules showed a weaker range of correlations when compared to transcripts covered by the 770 NanoString probes (Figure 5B). Notably, we observed an age specific effect in which the correlation between nCounter probe expression and RNA-Seq transcripts increased over time (Figure 5B). A mild correlation at around three months of age (Pearson’s correlation coefficient: 0.39) increased to a moderate correlation at 12 months of age (Pearson’s correlation coefficient: 0.51). Furthermore, we observed a high correlation of log count values for the majority of NanoString probes when compared to log TPM transcript ratios from RNA-Seq data. The majority of the 770 measured NanoString probes (716/770 probes, 93%) were positively correlated with RNA-Seq transcripts (Figure S2). In order to test whether noise introduced by highly variable transcripts affects the correlation between NanoString probes and RNA-Seq transcripts, Pearson’s correlation coefficients and variance in RNA-Seq expression across 137 samples were compared. There was no significant trend indicating an effect of highly variable transcripts on the overall correlation coefficients between transcripts measured by RNA-Seq and NanoString (Figure S2).

**Figure 5:**
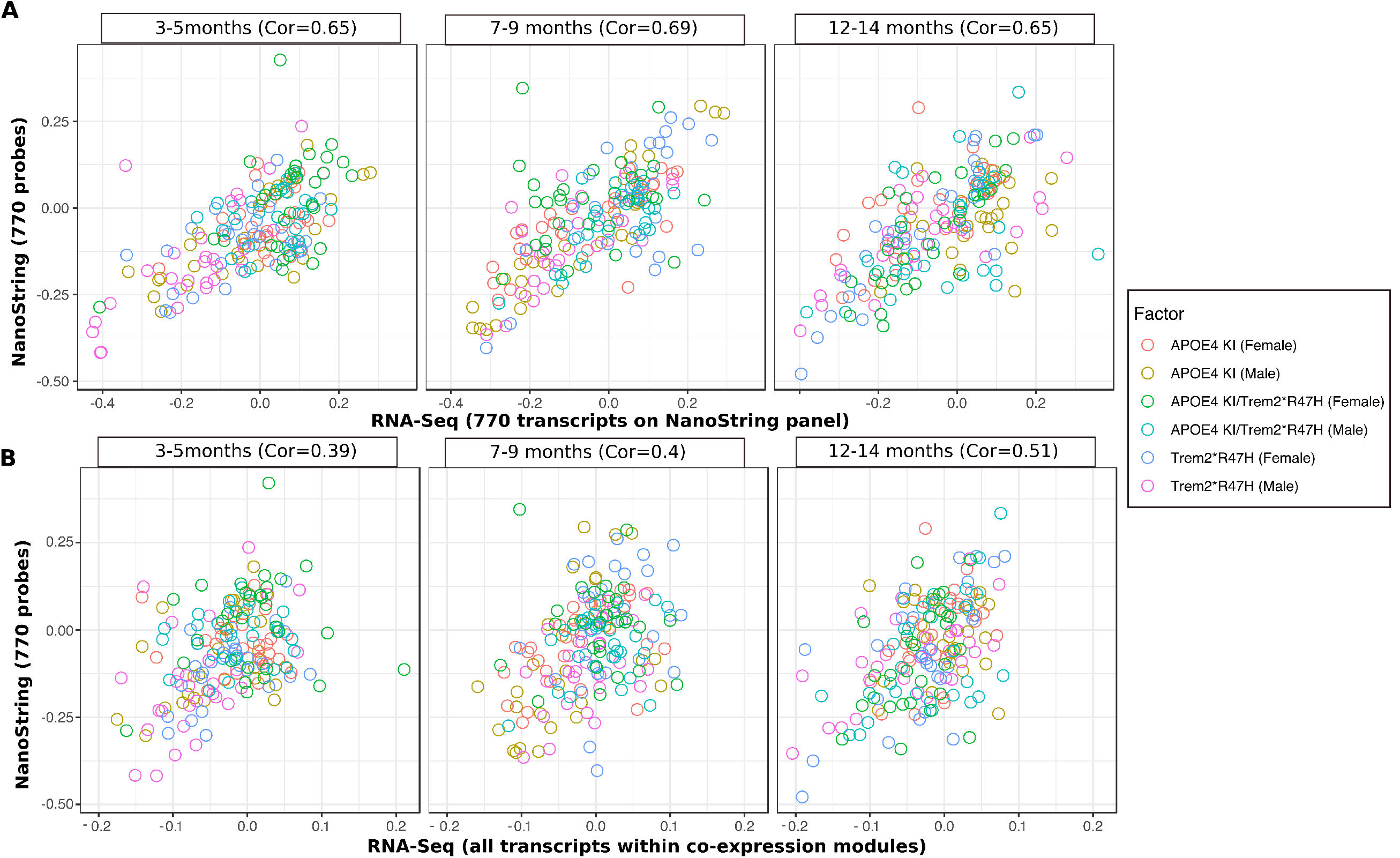
Platform comparison between the nCounter Mouse AD panel and RNA-Seq correlation with AMP-AD modules across 137 samples. The plots display the correlation between human AMP-AD co-expression modules and gene expression profiles derived from the NanoString panel and RNA-Seq data for the same 137 mouse samples. A detailed comparison is provided for three different age stages and three mouse models carrying LOAD associated risk variants. **(A)** A strong positive correlation was observed across all ages and samples combined when comparing expression of the 770 transcripts on the NanoString panel. **(B)** The correlation between NanoString and RNA-Seq expression analysis decreased overall when comparing all module transcripts measured by RNA-Seq to the subset of 770 probes on the NanoString panel. However, an age specific effect was observed for the mouse transcripts in which correlation with human co-expression modules increased with age.

## DISCUSSION

Here, we describe a novel systems biology approach to rapidly assess disease relevance for three novel mouse models carrying two human risk variants, strongly associated with LOAD. The nCounter Mouse AD gene expression panel was designed to align human brain transcriptome data covering 30 co-expression modules. Cross-species comparison of human and mouse revealed that immune associated co-expression modules which harbor genes that have recently diverged in sequence were more likely to be lowly expressed or absent at the transcript level in brains from 6 month old B6 mice. In contrast, neuronal modules containing genes with a lower degree of sequence divergence between both species were more likely to be highly and constitutively expressed in the mouse brain when compared to the remaining co-expression modules. This is in line with evidence from multiple studies highlighting that conserved neuronal process in the brain are under strong purifying selection while immune related genes are more likely to diverge in function and expression patterns across species (13,14). By using our prioritization approach, we selected for 760 key mouse genes targeting a subset of highly co-expressed human genes. This subset of genes on the NanoString panel showed overall lower levels of sequence divergence compared to human genes and higher expression levels in the mouse brain, reducing potential noise introduced by lowly expressed transcripts across expression modules.

Furthermore, we observed a robust and significant correlation between human co-expression modules and three mouse models carrying two LOAD associated risk variants (APOE4 KI, Trem2*R47H, APOE4 KI/Trem2*R47H). Cross-platform comparison between the novel Mouse AD panel and RNA-Seq data revealed a strong correlation between mouse gene expression changes independent of platform related effects. Notably, the correlation between nCounter probe and RNA-Seq transcript expression with human co-expression modules was highest in aged mice older than 12 months. This age-dependent overlap might be expected due to the late-onset nature of Alzheimer’s disease resulting in an increased number of highly co-expressed genes in aged mice carrying human LOAD risk variants. In addition, the strongest correlation between human and mouse module signatures was observed when using the subset of 770 transcripts on the NanoString panel. This highlights that assessment of key genes in the brain, contributing highly to module expression, can improve the characterization of novel LOAD mouse models and their alignment with specific human co-expression modules.

Interestingly, novel LOAD mouse models showed better concordance with distinct human co-expression modules, reflecting a different transcriptional response driven by the human *APOE* and *TREM2* associated LOAD risk variants. The strong negative correlation between the Trem2*R47H knock-in mice and immune related human co-expression highlights the important role of the LOAD associated *TREM2* R47H variant in Alzheimer’s related immune processes. This effect is reproducible across human co-expression modules, which derive from three independent cohorts and five different brain regions (cerebellum, frontal cortex, temporal gyrus, frontal gyrus, frontal pole). Similarly, a strong negative correlation between co-expression modules associated with cell cycle and DNA repair was observed for the mouse APOE4 KI model. This overlap with human late-onset co-expression signatures early in life was observed for a number of different brain regions and is absent in Trem2*R47H knock-in mice. Furthermore, aged APOE4 KI mice show a strong overlap with several human neuronal co-expression modules enriched for genes that play an important role in synaptic signaling and myelination. Although, APOE4 KI mice lack a clear neurodegenerative phenotype, this age dependent shift in co-expression patterns associated with core LOAD pathologies points to an increased susceptibility of cognitive decline in aged mice. This is in line with several studies, which have shown that cognitive deficits in APOE4 transgenic mice develop late in life (15,16).

### Limitations of the approach

Albeit being an excellent resource for characterizing molecular pathways and key drivers of disease, co-expression modules based on human post-mortem brain data have several limitations. They might not reflect changes that occur early in disease pathogenesis. In addition, although a high concordance was observed across brain regions for the 30 modules, they might not cover individual or region-specific differences in patients in response to amyloid and tau pathology (12). Furthermore, we used brain homogenates from our mouse models for the transcript comparison with different human brain regions in this study. Dissection of mouse brain regions to match the human studies might further improve the observed co-expression module correlations.

## CONCLUSIONS

Taken together, we show that the novel nCounter Mouse AD gene expression panel offers a rapid and cost-effective approach to assess disease relevance of novel LOAD mouse models. Furthermore, this co-expression based approach offers a high level of reproducibility and will supplement methods solely based on differential expression analysis. Ultimately, this will help us to better understand the relevance of novel LOAD mouse models in regard to specific pathways and processes contributing to late-onset Alzheimer’s disease.

## METHODS

### AMP-AD post-mortem brain cohorts and gene co-expression modules

Data on the 30 human AMP-AD co-expression modules was obtained from the Synapse data repository (DOI: 10.7303/syn11932957.1). The modules derive from three independent LOAD cohorts, including 700 samples from the ROSMAP cohort, 300 samples from the Mount Sinai Brain bank and 270 samples from the Mayo cohort. Details on post-mortem brain sample collection, tissue and RNA preparation, sequencing, and sample QC can be found in previously published work related to each cohort (10,11,17). A detailed description on how co-expression modules were identified can be found in the recent study that identified the harmonized human co-expression modules as part of transcriptome wide AD meta-analysis (12). Briefly, Logsdon et al. performed library normalization and covariate adjustments for each human study separately using fixed/mixed effects modeling to account for batch effects. Among the 2,978 AMP-AD modules identified across all tissues (10.7303/syn10309369.1), 660 modules were selected by Logsdon et al. which showed an enrichment for at least one AD-specific differential expressed gene set from the meta-analysis (10.7303/syn11914606) in cases compared to controls. Lastly, the edge betweenness graph clustering method was applied to identify 30 aggregate modules that are not only differentially expressed but are also replicated across multiple independent co-expression module algorithms (12). Among the 30 aggregate co-expression modules, five consensus clusters have been described by Logsdon et al. (12). These consensus clusters consist of a subset of modules which are associated with similar AD related changes across the multiple studies and brain regions. Here, we used Reactome pathway (https://reactome.org/) enrichment analysis to identify specific biological themes across these five consensus clusters. A hypergeometric model, implemented in the clusterProfiler R package (18), was used to assess whether the number of selected genes associated within each set of AMP-AD modules defining a consensus cluster was larger than expected. All p-values were calculated based the hypergeometric model (19). Pathways were ranked based on their Bonferroni corrected p-values to account for multiple testing. Finally, consensus clusters were annotated based on the highest ranked and non-overlapping term for each functionally distinct cluster.

### Selection of NanoString probes for the nCounter Mouse AD Panel

Since NanoString gene expression panels are comprised of 770 probes with the option to customize 30 additional probes, we developed a formal prioritization procedure to identify the most representative genes and ensure broadest coverage across all modules (Figure 1). Expression and transcript annotations for the 30 human co-expression modules were obtained via the AMP-AD knowledge portal (https://www.synapse.org/#!Synapse:syn11870970/tables/). To prioritize probe targets for the novel Mouse AD panel, human genes were ranked within each of the human AMP-AD co-expression modules based on their percentage of variation explaining the overall module behavior. First, we calculated a gene ranking score by multiplying correlations of transcripts with the percentage of variation explained by the first five principal components within each of the aggregated human AMP-AD modules. Secondly, the sums of the resulting gene scores for the first five principal components were calculated and converted to absolute values in order to rank highly positive or negative correlated transcripts within each human co-expression module. As a next step, only human transcripts with corresponding one-to-one mouse orthologous genes that are expressed in whole-brain tissue samples from six-month-old B6 mice were retained for downstream prioritization. Furthermore, we included information on drug targets for LOAD from the AMP-AD Agora platform (agora.ampadportal.org), as nominated by members of the AMP-AD consortium (10.7303/syn2580853). A total of 30 AMP-AD drug discovery targets that were highly ranked in our gene ranking approach and nominated by multiple AMP-AD groups were included on the panel (Supplemental Table 3). Finally, ten housekeeping genes (*AARS*, *ASB7*, *CCDC127*, *CNOT10*, *CSNK2A2*, *FAM104A*, *LARS*, *MTO1*, *SUPT7L*, *TADA2B*) were included on the panel as internal standard references for probe normalization. This resulted in a total of 770 proposed NanoString probes, targeting the top 5% of ranked genes for each human AMP-AD expression module.

### nCounter Mouse AD Panel Probe Design

The probe design process breaks a transcript’s sequence down into 100 nucleotide (nt) windows to profile for probe characteristics, with the final goal of choosing the optimal pair of adjacent probes to profile any given target. Each window is profiled for intrinsic sequence makeup – non-canonical bases, G/C content, inverted and direct repeat regions, runs of poly-nucleotides, as well as the predicted melting temperature (Tm) for each potential probe-to-target interaction. The window is then divided in half to generate a probe pair, wherein each probe is thermodynamically tuned to determine the optimal probe length (ranging in size from 35-50 nt) within the 100 nt target region. Next, a cross-hybridization score is calculated for each probe region, using BLAST (20) to identify potential off-target interactions. In addition to a cross-hybridization score, a splice isoform coverage score was generated to identify transcripts that are isoforms of the gene intended to be targeted by the probe in question. Once all of this information is compiled, the final probe is then selected by identifying the candidate with the optimal splice form coverage, cross-hybridization score, and thermodynamic profile.

### In-silico panel QC for intramolecular interactions

To ensure that there are no potential intramolecular probe-probe interactions that could cause elevated background for any individual probe pair, a stringent intermolecular screen is run on every collection of probes assembled into a panel. A sensitive algorithm was used that calculates both the Tm and the free energy potential of interactions between every possible pair of probes in the project. If two probes conflict in a way that would likely cause background based on this calculation, an alternative probe is selected for one of the targets and the screening is re-run until there are no known conflicts.

### Mouse models

All experiments involving mice (Supplemental Table S4) were conducted in accordance with policies and procedures described in the Guide for the Care and Use of Laboratory Animals of the National Institutes of Health and were approved by the Institutional Animal Care and Use Committee at The Jackson Laboratory. All mice were bred and housed in a 12/12 hour light/dark cycle. All experiments were performed on a unified genetic background (C57BL/6J).

### Mouse brain sample collection

Upon arrival at the terminal endpoint for each aged mouse cohort, individual animals were weighed prior to intraperitoneal administration of ketamine (100mg/kg) and xylazine (10mg/kg). First confirming deep anesthetization via toe pinch, an incision was made along the midline to expose the thorax and abdomen followed by removal of the lateral borders of the diaphragm and ribcage revealed the heart. A small cut was placed in the right atrium to relieve pressure from the vascular system before transcardially perfusing the animal with 1XPBS via injection into the left ventricle. With the vascular system cleared, the entire brain was carefully removed and weighed before hemisecting along the midsagittal plane. Hemispheres were immediately placed in a cryovial and snap-frozen on dry ice. Brain samples were stored at −80°C until RNA extraction was performed.

### RNA sample preparation

RNA was isolated from tissue using the MagMAX mirVana Total RNA Isolation Kit (ThermoFisher) and the KingFisher Flex purification system (ThermoFisher, Waltham, MA). Brain hemispheres were thawed to 0°C and were lysed and homogenized in TRIzol Reagent (ThermoFisher). After the addition of chloroform, the RNA-containing aqueous layer was removed for RNA isolation according to the manufacturer’s protocol, beginning with the RNA bead binding step. RNA concentration and quality were assessed using the Nanodrop 2000 spectrophotometer (Thermo Scientific) and the RNA Total RNA Nano assay (Agilent Technologies, Santa Clara, CA).

### RNAseq library preparation and data collection

Sequencing libraries were constructed using TruSeq DNA V2 (Illumina, San Diego, CA) sample prep kits and quantified using qPCR (Kapa Biosystems, Wilmington, MA). The mRNA was fragmented, and double-stranded cDNA was generated by random priming. The ends of the fragmented DNA were converted into phosphorylated blunt ends. An ‘A’ base was added to the 3’ ends. Illumina®-specific adaptors were ligated to the DNA fragments. Using magnetic bead technology, the ligated fragments were size-selected and then a final PCR was performed to enrich the adapter-modified DNA fragments, since only the DNA fragments with adaptors at both ends will amplify. Libraries were pooled and sequenced by the Genome Technologies core facility at The Jackson Laboratory. Samples were sequenced on Illumina HiSeq 4000 using HiSeq 3000/4000 SBS Kit reagents (Illumina), targeting 30 million read pairs per sample. Samples were split across multiple lanes when being run on the Illumina HiSeq, once the data was received the samples were concatenated to have a single file for paired-end analysis.

### NanoString gene expression panel and data collection

The NanoString Mouse AD gene expression panel was used for gene expression profiling on the nCounter platform (NanoString, Seattle, WA) as described by the manufacturer. nSolver software was used for analysis of NanoString gene expression values.

### Normalization of NanoString data

Normalization was done by dividing counts within a lane by geometric mean of the housekeeping genes from the same lane. For the downstream analysis, counts were log-transformed from normalized count values.

### Mouse-human expression comparison

First, we performed differential gene expression analysis for each mouse model and sex using the voom-limma (21) package in R. Secondly, we computed correlation between changes in expression (log fold change) for each gene in a given module with each mouse model, sex and age. Correlation coefficients were computed using cor.test function built in R as:

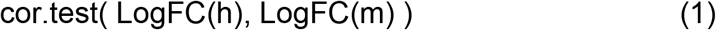

where LogFC(h) is the log fold change in transcript expression of human AD patients compared to control patients and LogFC(m) is the log fold change in expression of mouse transcripts compare to control mouse models. LogFC values for human transcripts were obtained via the AMP-AD knowledge portal (https://www.synapse.org/#!Synapse:syn11180450).

### Quality control of RNA-Seq data and read alignment

Sequence quality of reads was assessed using FastQC (v0.11.3, Babraham). Low-quality bases were trimmed from sequencing reads using Trimmomatic (v0.33) (22). After trimming, reads of length longer than 36 bases were retained. The average quality score at each base position was greater than 30 and sequencing depth were in range of 60 – 120 million reads. All RNA-Seq samples were mapped to the mouse genome (mm10 reference, build 38, ENSEMBL) using ultrafast RNA-Seq aligner STAR (23) (v2.5.3). The genes annotated for mm10 (GRCm38) were quantified in two ways: Transcripts per million (TPM) using RSEM (v1.2.31) and raw read counts using HTSeq-count (v0.8.0).

### Mouse-human co-expression module conservation

Genomic information on orthologous groups was obtained via the latest ENSEMBL build for human genome version GRCh38. All orthologous relationships were downloaded via BioMart (24) (biomart.org). dN/dS statistics were retrieved for all orthologous gene pairs with a one-to-one relationship between human and mouse. dN/dS values are calculated as the ratio of nonsynonymous substitutions to the number of synonymous substitutions in protein coding genes. The dN/dS values in ENSEMBL were calculated based on the latest version of the codeml (http://abacus.gene.ucl.ac.uk/software/paml.html) package using standard parameters (ensembl.org/info/genome/compara/homology_method.html) (25).

## DECLARATIONS

## Acknowledgments

We thank the many institutions and their staff that provided support for this study and who were involved in this collaboration. We would like to acknowledge Jamie Kuhar for her critically reading of the manuscript.

## Availability of data and materials

The results published here are in whole or in part based on data obtained from the AMP-AD Knowledge Portal (doi:10.7303/syn2580853). ROSMAP Study data were provided by the Rush Alzheimer’s Disease Center, Rush University Medical Center, Chicago. Data collection was supported through funding by NIA grants P30AG10161, R01AG15819, R01AG17917, R01AG30146, R01AG36836, U01AG32984, U01AG46152, the Illinois Department of Public Health, and the Translational Genomics Research Institute. Mayo RNAseq Study data were provided by the following sources: The Mayo ClinicAlzheimer’s Disease Genetic Studies, led by Dr. Nilufer Ertekin-Taner and Dr. Steven G. Younkin, Mayo Clinic, Jacksonville, FL using samples from the Mayo Clinic Study of Aging, the Mayo Clinic Alzheimer’s Disease Research Center, and the Mayo Clinic Brain Bank. Data collection was supported through funding by NIA grants P50 AG016574, R01 AG032990, U01 AG046139, R01 AG018023, U01 AG006576, U01 AG006786, R01 AG025711, R01 AG017216, R01 AG003949, NINDS grant R01 NS080820, CurePSP Foundation, and support from Mayo Foundation. Study data includes samples collected through the Sun Health Research Institute Brain and Body Donation Program of Sun City, Arizona. The Brain and Body Donation Program is supported by the National Institute of Neurological Disorders and Stroke (U24 NS072026 National Brain and Tissue Resource for Parkinson’s Disease and Related Disorders), the National Institute on Aging (P30 AG19610 Arizona Alzheimer’s Disease CoreCenter), the Arizona Department of Health Services (contract 211002, Arizona Alzheimer’s Research Center), the Arizona Biomedical Research Commission (contracts 4001, 0011, 05-901 and 1001 to the Arizona Parkinson’s Disease Consortium) and the Michael J. Fox Foundation for Parkinson’s Research. MSBB data were generated from postmortem brain tissue collected through the Mount Sinai VA MedicalCenter Brain Bank and were provided by Dr. Eric Schadt from Mount Sinai School of Medicine. Mouse RNAseq data from the MODEL-AD consortium is available through Synapse via the AMP-AD knowledge portal (www.synapse.org/#!Synapse:syn17095980)

## Authors’ contributions

CP designed the novel transcriptome panel and performed bioinformatics analyses. RP, AF, AU, TP performed the gene-expression analyses in human and mouse brain tissue. EP designed the NanoString probes and guided the creation of the novel NanoString panel. BAL and LM curated human brain data. DG, GRH and MS performed mouse experiments. GWC and MS supervised and designed the project. All authors read and approved the manuscript. CP, GWC and RP wrote the manuscript.

## Ethics approval

All experiments involving mice were conducted in accordance with policies and procedures described in the Guide for the Care and Use of Laboratory Animals of the National Institutes of Health and were approved by the Institutional Animal Care and Use Committee at The Jackson Laboratory.

## Consent for publication

All authors have approved of the manuscript and agree with its submission.

