## Supplemental Materials (Figures and Tables) for "A novel systems biology approach to evaluate mouse models of late-onset Alzheimer’s disease"

### SUPPLEMENTAL MATERIAL

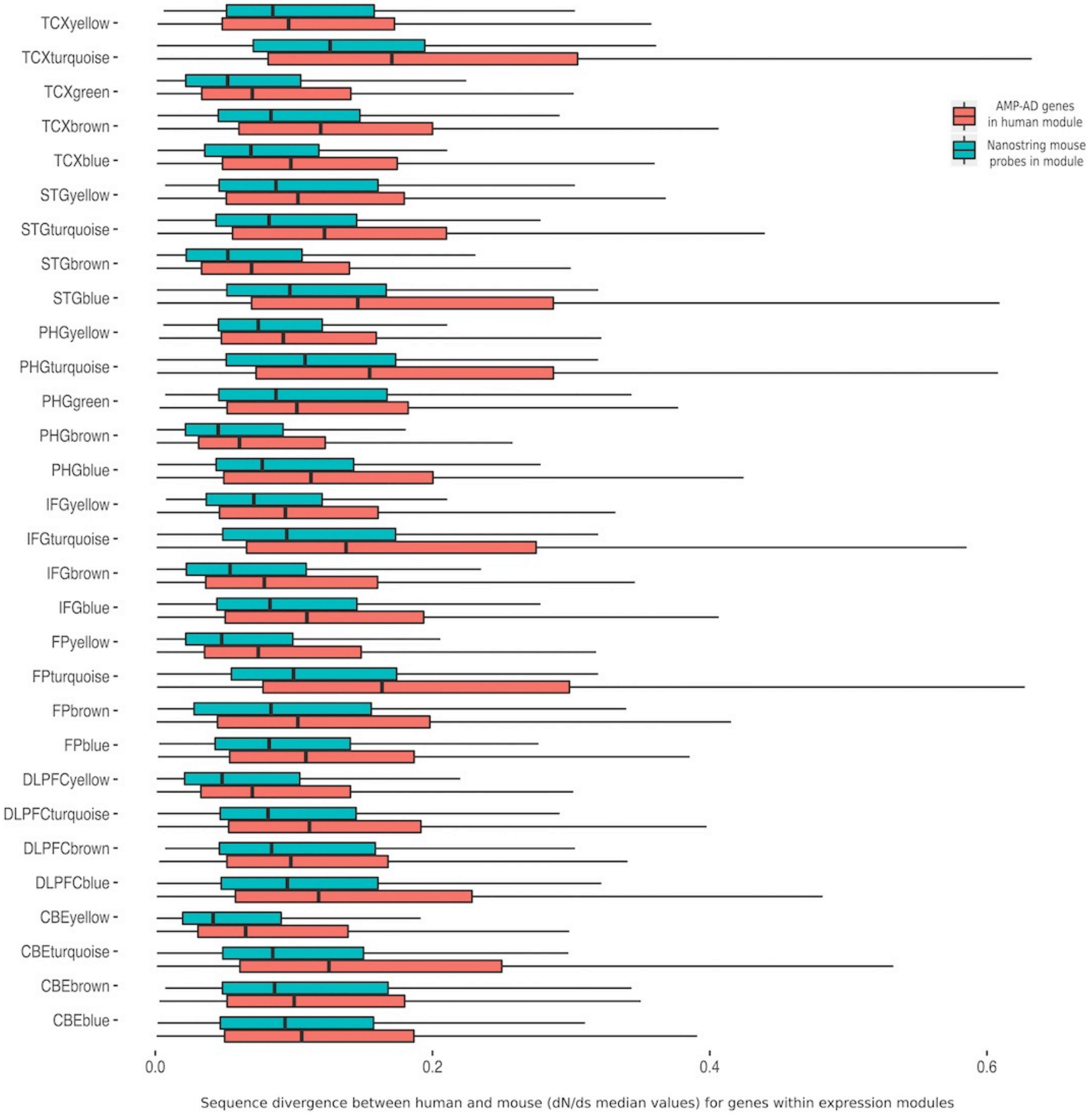


**Supplemental Figure 1: Sequence divergence of genes prioritized for the nCounter Mouse AD panel compared to all genes in each AMP-AD module**

For each of the 30 human co-expression modules, selected key genes on the Mouse AD panel are more conserved when compared the overall gene content.

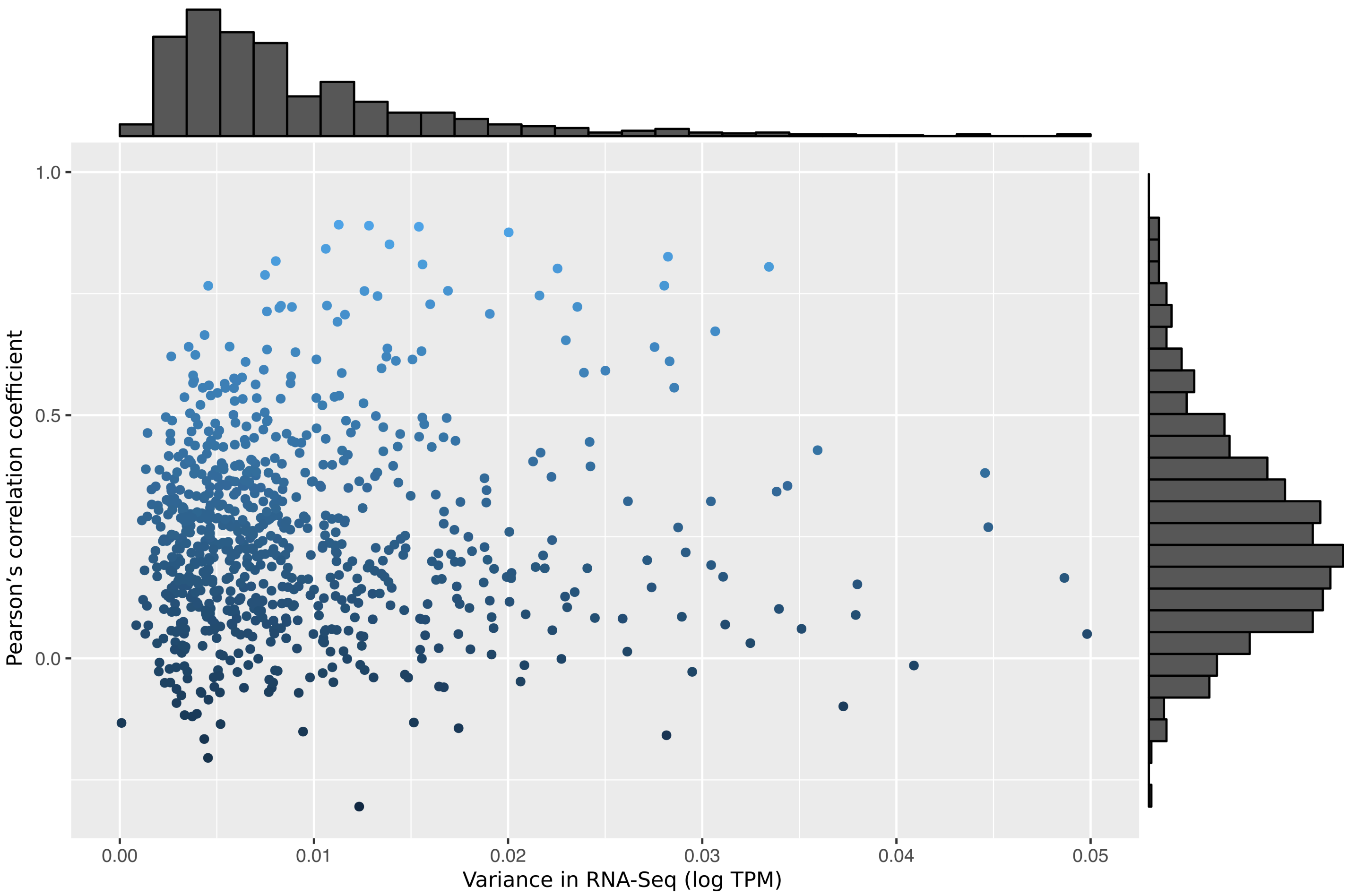


**Supplemental Figure 2: Comparison between Pearson's correlation coefficients and variance in RNA-Seq expression for each probe across 137 samples**

The scatter plot depicts the correlation between the variance of each transcript measured by RNA-Seq versus the Pearson's correlation coefficient between the NanoString and RNA-Seq expression for each probe across 137 samples. Of the 770 probes, 492 showed significant positive correlation (Pearson's correlation coefficient > 0.17, p-value < 0.05), while 223 showed positive but insignificant correlation (0 < Pearson's correlation coefficient < 0.17, p-value > 0). Two probes showed significant negative correlation (Pearson's correlation coefficient < -0.20, p-value < 0) and 61 showed negative insignificant correlation (-0.30 < Pearson's correlation coefficient < 0, p-value > 0). We did not observe any significant trend between variance and correlation coefficients.

**SUPPLEMENT TABLE 1: Reactome pathway annotation for the five consensus clusters**

The table depicts the Reactome pathway annotations for the five functional consensus clusters associated with the 30 human co-expression modules. The highest ranked and non-overlapping Reactome pathway annotations are highlighted and were used to annotate the five consensus clusters (A-E).

| Consensus Cluster | Description | GeneRatio | adjusted p-value | qvalue |
|---|---|---|---|---|
| A | **Extracellular matrix organization** | 64/834 | 1.50E-06 | 1.39E-06 |
| A | Diseases associated with glycosaminoglycan metabolism | 16/834 | 1.50E-06 | 1.39E-06 |
| A | Diseases of glycosylation | 16/834 | 1.50E-06 | 1.39E-06 |
| A | Defective B4GALT7 causes EDS, progeroid type | 13/834 | 4.16E-06 | 3.88E-06 |
| A | Defective B3GAT3 causes JDSSDHD | 13/834 | 4.16E-06 | 3.88E-06 |
| B | **Immune System** | 311/1517 | 2.29E-13 | 1.97E-13 |
| B | Cytokine Signaling in Immune system | 123/1517 | 3.32E-13 | 2.87E-13 |
| B | Extracellular matrix organization | 113/1517 | 1.73E-13 | 1.49E-13 |
| B | Interferon gamma signaling | 51/1517 | 9.92E-14 | 8.55E-14 |
| B | Non-integrin membrane-ECM interactions | 38/1517 | 6.22E-12 | 5.36E-12 |
| C | **Neuronal System** | 178/2046 | 1.59E-30 | 1.30E-30 |
| C | Transmission across Chemical Synapses | 122/2046 | 1.22E-18 | 1.00E-18 |
| C | Neurotransmitter Receptor Binding And Downstream Transmission In The Postsynaptic Cell | 87/2046 | 2.28E-13 | 1.87E-13 |
| C | Potassium Channels | 65/2046 | 1.47E-10 | 1.20E-10 |
| C | Voltage gated Potassium channels | 37/2046 | 5.47E-11 | 4.47E-11 |
| D | Gene Expression | 271/1614 | 1.24E-05 | 1.06E-05 |
| D | **Cell Cycle, Mitotic** | 142/1614 | 5.34E-05 | 4.57E-05 |
| D | Nonsense-Mediated Decay (NMD) | 52/1614 | 2.49E-05 | 2.13E-05 |
| D | Nonsense Mediated Decay (NMD) enhanced by the Exon Junction Complex (EJC) | 52/1614 | 2.49E-05 | 2.13E-05 |
| D | Influenza Viral RNA Transcription and Replication | 52/1614 | 5.34E-05 | 4.57E-05 |
| E | Gene Expression | 517/2390 | 1.46E-55 | 1.01E-55 |
| E | **Organelle biogenesis and maintenance** | 178/2390 | 4.15E-19 | 2.89E-19 |

**SUPPLEMENT TABLE 2: Summary statistics nCoutner Mouse AD panel**
Summary statistics for the 30 human co-expression modules, including the annotation for brain regions and cohorts from which the specific modules were generated. In addition, NanoString probe coverage and functional consensus cluster membership are listed for each co-expression module.

| Genes in AMP-AD module | Covered Nanostring probes | Coverage [%] | Name | Brain Region | Cohort | Consensus Cluster |
|---|---|---|---|---|---|---|
| 1713 | 119 | 7 | TCXblue | temporal cortex | Mayo | A |
| 743 | 84 | 11 | IFGyellow | inferiorfrontal gyrus | Mount Sinai Brain Bank | A |
| 910 | 101 | 11 | PHGyellow | parahipocampal gyrus | Mount Sinai Brain Bank | A |
| 1751 | 183 | 10 | DLPFCblue | dorsolateral prefrontal cortex | ROS/MAP | B |
| 1977 | 200 | 10 | CBEturquoise | cerebellum | Mayo | B |
| 1131 | 78 | 7 | TCXturquoise | temporal cortex | Mayo | B |
| 1456 | 126 | 9 | IFGturquoise | inferiorfrontal gyrus | Mount Sinai Brain Bank | B |
| 1171 | 143 | 12 | STGblue | superiortemporal gyrus | Mount Sinai Brain Bank | B |
| 1195 | 104 | 9 | PHGturquoise | parahipocampal gyrus | Mount Sinai Brain Bank | B |
| 1001 | 107 | 11 | FPturquoise | frontal pole | Mount Sinai Brain bank | B |
| 3019 | 192 | 6 | DLPFCyellow | dorsolateral prefrontal cortex | ROS/MAP | C |
| 1739 | 157 | 9 | CBEyellow | cerebellum | Mayo | C |
| 2766 | 186 | 7 | TCXgreen | temporal cortex | Mayo | C |
| 4673 | 234 | 5 | IFGbrown | inferiorfrontal gyrus | Mount Sinai Brain Bank | C |
| 3414 | 211 | 6 | STGbrown | superiortemporal gyrus | Mount Sinai Brain Bank | C |
| 2123 | 165 | 8 | PHGbrown | parahipocampal gyrus | Mount Sinai Brain Bank | C |
| 4426 | 188 | 4 | FPyellow | frontal pole | Mount Sinai Brain Bank | C |
| 882 | 139 | 16 | DLPFCbrown | dorsolateral Prefrontal Cortex | ROS/MAP | D |
| 504 | 95 | 19 | CBEbrown | cerebellum | Mayo | D |
| 2013 | 151 | 8 | TCXyellow | temporal cortex | Mayo | D |

| | | | | | | |
|---|---|---|---|---|---|---|
| 2885 | 236 | 8 | IFGblue | inferiorfrontal gyrus | Mount Sinai Brain Bank | D |
| 1799 | 159 | 9 | STGyellow | superiortemporal gyrus | Mount Sinai Brain Bank | D |
| 1151 | 139 | 12 | PHGgreen | parahipocampal gyrus | Mount Sinai Brain Bank | D |
| 1991 | 278 | 14 | FPblue | frontal pole | Mount Sinai Brain Bank | D |
| 2489 | 144 | 6 | DLPFCturquoise | dorsolateral prefrontal cortex | ROS/MAP | E |
| 4509 | 177 | 4 | CBEblue | cerebellum | Mayo | E |
| 1851 | 125 | 7 | TCXbrown | temporal cortex | Mayo | E |
| 2404 | 119 | 5 | STGturquoise | superiortemporal gyrus | Mount Sinai Brain Bank | E |
| 3733 | 170 | 5 | PHGblue | parahipocampal gyrus | Mount Sinai Brain Bank | E |
| 1289 | 76 | 6 | FPbrown | frontal pole | Mount Sinai Brain Bank | E |

**SUPPLEMENT TABLE 3: Annotations for selected NanoString probes covering human co-expression modules**

The attached table contains human gene to mouse probe annotations for the 770 selected probes on the nCounter Mouse AD panel. Individual mouse probes were assigned to multiple human co-expression modules derived from different brain regions or cohorts. Gene scores indicate contribution of each probe set to relative module behavior for each assigned human co-expression module. AMP-AD drug targets as listed by Agora (agora.ampadportal.org) and housekeeping genes are highlighted (**See attached Excel table**).

**SUPPLEMENT TABLE 4: Overview of mouse samples used for strain survey**

| Mouse model | Strain Nomenclature (Number) | Total Samples | Sex | 3-5 months | 7-9 months | 12-14 months |
|---|---|---|---|---|---|---|
| B6 | C57BL/6J (JAX#664) | 35 | Female | 6 | 6 | 5 |
| | | | Male | 5 | 6 | 7 |
| APOE4 KI | B6.Cg-Apoe$^{tm1.1(APOE*4)Adiuj}$/J (JAX # 27894) | 32 | Female | 6 | 6 | 5 |
| | | | Male | 5 | 6 | 4 |
| Trem2*R47H | B6.Cg-Apoe$^{tm1.1(APOE*4)Adiuj}$Trem2$^{em1Adiuj}$/J (JAX #27918) | 35 | Female | 5 | 6 | 6 |
| | | | Male | 6 | 6 | 6 |
| APOE4 KI/Trem2*R47H | B6.Cg-Apoe$^{tm1.1(APOE*4)Adiuj}$Trem2$^{em1Adiuj}$/J (JAX #28709) | 35 | Female | 6 | 6 | 5 |
| | | | Male | 5 | 6 | 7 |
